# The Findings Workflow: An agentic, collaborative, provenance-tracked framework for scientific data analysis with Claude Code

**DOI:** 10.64898/2026.10.28.762630

**Authors:** Michael Riffle, Alex Zelter, Michael J MacCoss

## Abstract

Scientists presented with quantitative data often face two related problems: confidence that an analysis is statistically and computationally correct, and provenance sufficient to reproduce the resulting figures and conclusions. Large language models (LLMs) can make sophisticated analyses accessible through a collaborative, conversational interface, but unstructured use can worsen both problems by producing plausible but incorrect results and obscuring the steps that produced them. Here we present the Findings Workflow, a free, open-source Claude Code plugin designed to retain the flexibility of conversational LLM-assisted analysis while placing it within a structured, verifiable, and reproducible environment. The workflow uses staged, gated progression to ensure the science, metadata, and data are understood and tested data loaders are built before downstream analysis begins. Specialized subagents perform statistical analysis, programming, visualization, research, reporting, and validation, with independent review incorporated throughout. The plugin also provides vetted analysis templates, conventions, and methodological guidance that serve as trusted sources of opinion when deciding how analyses should be performed. Scientists remain actively involved throughout, exploring their data and directing the analysis through natural-language conversation, while the workflow supplies structure, verification, and persistent state around that collaboration. Findings are recorded automatically together with the data, code, methods, figures, and provenance needed to reproduce them. We demonstrate the workflow with a worked example that reproduces central findings from a published plasma extracellular vesicle proteomics dataset examining Alzheimer’s disease (Mag-Net). Together, these features are designed to make LLM-assisted scientific analysis easier to trust, inspect, and reproduce without sacrificing its conversational and exploratory nature.

## Introduction

The final output of raw quantitative proteomics analysis is typically a matrix of samples and features (e.g., precursors, protein groups, or genes) with some measured value of abundance for those features in those samples. This matrix, along with metadata, is then analyzed downstream by a researcher for exploratory analysis and to confirm and test hypotheses. Standard examples of this type of analyses include biomarker discovery via feature finding or differential abundance analysis, developing machine learning classifiers to predict treatment or disease state, or regression models that can predict the level of exposure to a drug or environmental toxin, or even disease progression. However, this final statistical analysis can be a daunting task for many scientists and biomedical researchers who lack expertise in statistics or programming. While many excellent libraries exist for performing canonical, standard types of quantitative analysis, many of these are implemented in R^1^ and python^2^, and may lead to a lack of confidence that the analysis is being implemented correctly. Were the data read in correctly? Is the correct statistical test being performed? What assumptions do those tests have about the data and are they true? What options are there for normalization, batch correction, and missingness imputation? The simpler an analysis tool, the fewer options the researcher has and the more assumptions there are about the data. The more flexible and powerful a tool, the harder and more complicated it is to use with a larger chance for error. The ideal solution would be flexible, transparent, interactive, and easy to use.

Large language models (LLMs) bring an unprecedented amount of power to researchers looking to analyze their own data. Using a simple chat interface, researchers can (ostensibly) analyze their data using both canonical methods (for standard analysis) and bespoke methods for customized analysis answering their specific questions about their own data. Leveraging all of the statistical, programming, and science knowledge encoded in the LLM, they can rapidly produce figures and conclusions from their data that seem quite reasonable. However, it is well known that LLMs can (and often do) make mistakes. A researcher knowledgeable of this has not solved the confidence issue associated with doing the analysis themself–they have only made mistakes harder to find. Buried in the dense output of reasonable-looking results could be a faulty implementation of cross-validation, an error in loading the data or mapping it to metadata, or some level of imputation that went unnoticed by the researcher. Additionally, analysis with LLMs introduces a serious issue of provenance. After long chat sessions arrive at figures or conclusions the researcher thinks look reasonable, exactly how did they arrive there? Exactly how can that figure be reproduced? What were the exact analyses that led to a particular conclusion? Using LLMs for analysis in an unstructured, unverified way, only exacerbates issues of confidence and introduces issues of provenance.

Here we introduce the Findings Workflow, a Claude Code^3^ plugin designed specifically to allow researchers to confidently leverage LLMs for scientific data analysis. Although some LLM-based “chat with your data” tools exist, these are often simple, do not emphasize validation, or are locked into proprietary, commercial platforms^4,5^. These do little to address confidence and provenance. The Findings Workflow is built using principled rules that emphasize independently verifying all research and analysis and independently reviewing all outputs. Additionally, the findings workflow automatically records findings with full provenance as discoveries are made during conversations between the researcher and Claude. These findings are saved in structured markdown documents in a growing knowledgebase and include metadata as machine-readable frontmatter (e.g. tags, ontologies, keywords), and human-readable descriptions, discussions, figures, tables, and full, specific provenance (e.g., which scripts lead to which conclusions, generated which figures, and so on). As a Claude Code plugin, the Findings Workflow runs on a local computer and is simple to install, update, and use. All interaction is conversational. The Findings Workflow is built on a principled flow for data analysis, where the project and its goals must be understood first, then the metadata, then the data, then robust, highly-tested, and trusted data loaders written before any data analysis. Such rules are encoded as hard gates, ensuring no analysis is done before the data are understood. The Findings Workflow ships with dedicated agentic subsystems for different aspects of analysis, such as research, coding, statistics, and figure generation. It ships with example templates for implementing many types of canonical data analysis, quality control, and data visualization that are used as a basis of opinion for how analyses should be performed by agents on end user systems.

The Findings Workflow is open source and freely available at https://github.com/mriffle/findings-ai-collab-workflow.

## Implementation

### Claude Code plugin architecture

The Findings Workflow is implemented as a Claude Code plugin. Claude Code is an agentic harness that provides an environment for Claude (LLM models from Anthropic) to interact with a project. It allows Claude to directly read and write files on your computer, execute programs, and delegate work to specialized agents. Claude Code can run on the command line in a terminal, through integrated developer environments (IDEs), as a desktop application, or via the web. A Claude Code plugin is an installable package that extends this environment with reusable components such as skills that describe how particular tasks should be performed, specialized subagent definitions that run with specialized prompts in their own context, commands that define workflow entrypoints, event-driven hooks that can deterministically gate or block actions, example code, configurations, and (where appropriate) external tool integrations. All the components are installed and updated using a simple command in Claude Code, pulling the plugin files from GitHub^6^.

### Findings Workflow plugin architecture

The Findings Workflow plugin is designed as a reusable data analysis environment, where the project file system directory holds the actual study. The plugin contains the workflow logic, agents, procedures, conventions, enforcement mechanisms, and vetted data analysis and data visualization implementations. A project directory contains the researcher’s metadata, quantitative data (e.g., abundance matrices), and the temporary and permanent artifacts resulting from a given project’s analysis. These include learned descriptions of the project’s domain and purpose, descriptions of metadata and data, project state, adapted scripts, results, figures, research, findings, and reports. The plugin, itself, contains no study-specific data–only the framework and architecture for performing the analysis of any dataset. (Figure 1A)

**Figure 1.**
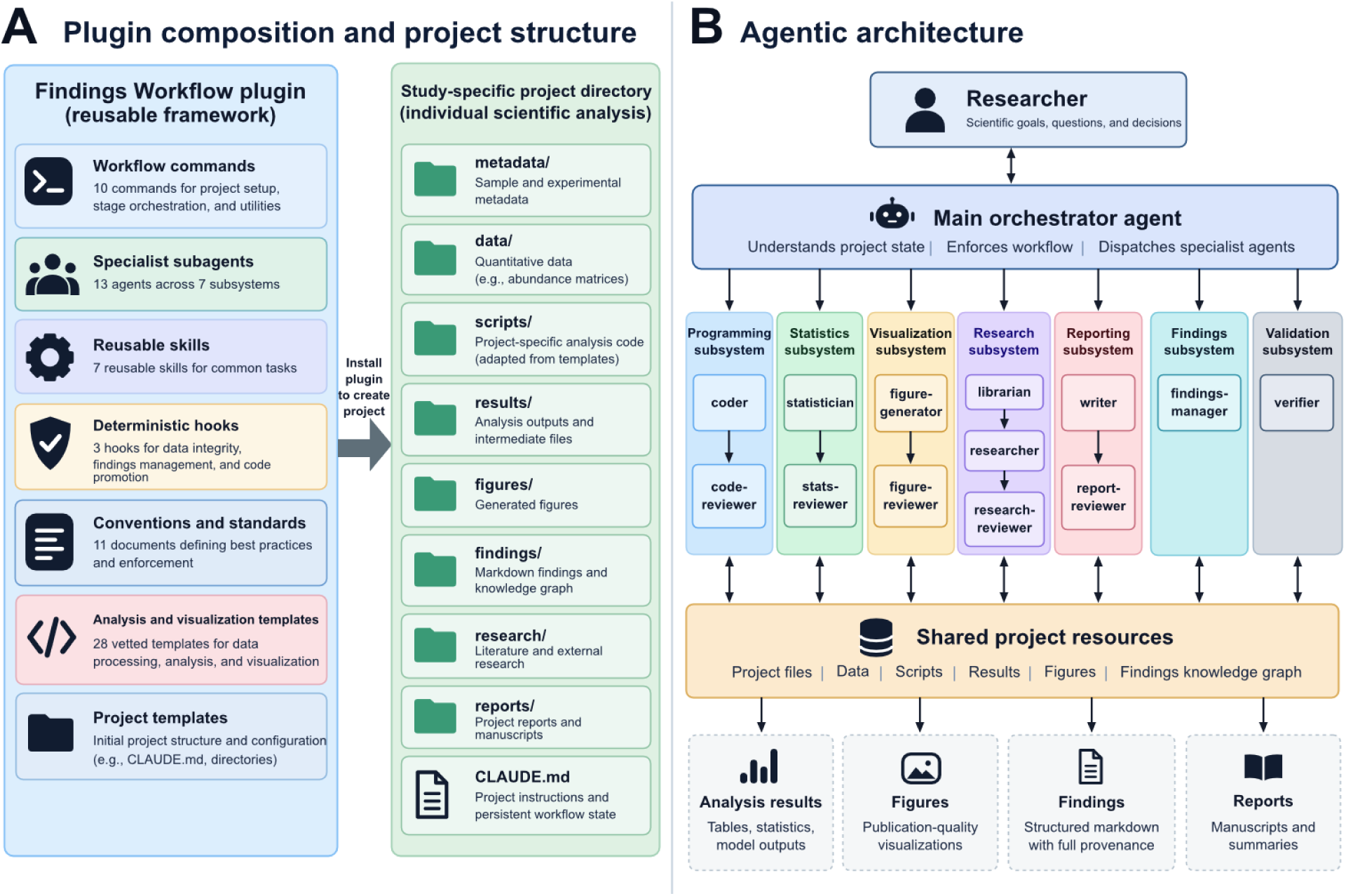
Overview of Findings Workflow architecture. (A) Plugin composition and project structure. The blue shaded section depicts the types of files and definitions shipped with the Findings Workflow Claude Code plugin. It contains workflow commands, specialist subagents, reusable skills, deterministic enforcement hooks, conventions and standards, analysis and visualization templates, and project templates. Installation creates a study-specific project directory structure (green shaded area) that contains the researcher’s data, analysis artifacts, findings, and reports. (B) Agentic architecture. The researcher interacts directly with the main orchestrator agent that spawns subagents dedicated to specific aspects of the analysis workflow. These agents are organized into seven subsystems (see Table 1), including programming, statistics, visualization, research, reporting, managing findings, and validation. All agents interact with shared project resources, including data, code, results, and the findings knowledge graph.

**Table 1.** The 13 subagents, organized by subsystem.

| Subsystem | Agents | Principal responsibility |
|---|---|---|
| Findings | findings-manager | Creation and maintenance of the findings knowledge graph. Does not decide the truth of findings, custodial only. |
| Validation | verifier | Blind, context-isolated independent re-derivation of findings. |
| Programming | coder, code-reviewer | Generate and fix analysis code and independently review it. |
| Statistics | statistician, stats-reviewer | Perform statistical analyses and independently review statistical appropriateness. |
| Visualization | figure-generator, figure-reviewer | Generate figures and review the generated output. |
| Research | librarian, researcher, research-reviewer | Manage external research, conduct external research, factually verify research and reference accuracy. |
| Reporting | writer, report-reviewer | Write findings into a report and verify claims are supported. |

#### Workflow Commands

In the Findings Workflow, commands define workflow orchestration rather than running analysis programs. They define what must happen at any particular stage, inspect (and enforce) the project state for prerequisites, dispatch specialized agents, and update the persistent workflow state. The Findings Workflow provides 10 user-facing commands for moving through the stages of data analysis (Figure 2). These include three utility commands: (1) `init`: Scaffolds the project by setting up folders, project instructions, and files to hold project state. This is run first, after the user has created the project directory and placed the data and metadata into a sub directory named “data”. (2) `setup-env`: Sets up the project-level Python environment, installing python locally in this directory if it is not found. (3) `status`: Shows where the project stands. Additionally, there are seven stage commands corresponding to steps in the scientific data analysis workflow. The order of these commands is enforced by built in gates, where some stages require completion of previous stages. The guiding principle is always to understand the underlying data before building on it. These commands are: (1) `stage0-science`: Claude interviews the scientist about the study: the domain, the design, what is being studied and why. (2) `stage1-metadata`: Claude works to deeply understand the metadata. It tests relationships that should hold and looks for confounds and imbalances. (3) `stage2-data`: Claude works to deeply understand the data: the matrix orientation, whether the data have been transformed or normalized, evaluate missingness, contaminants, or duplicates. (4) `stage3-loaders:` Claude builds and rigorously verifies tested loaders. It then performs an extensive evaluation of quality control statistics and generates visualizations and a report. (5) `stage4-explore`: This step is the heart of the workflow. Once the project, metadata, and data are fully understood and robustly being loaded, true analysis of the project data can begin. The user explores the data with Claude, whether performing canonical or bespoke analysis. Findings are automatically saved. (6) `stage5-validate`: Before a finding is considered “validated”, they are blindly and independently re-derived using a pre-specified verification criterion. (7) `stage6-report`: Compiles the researcher’s findings into a full, vetted report.

**Figure 2.**
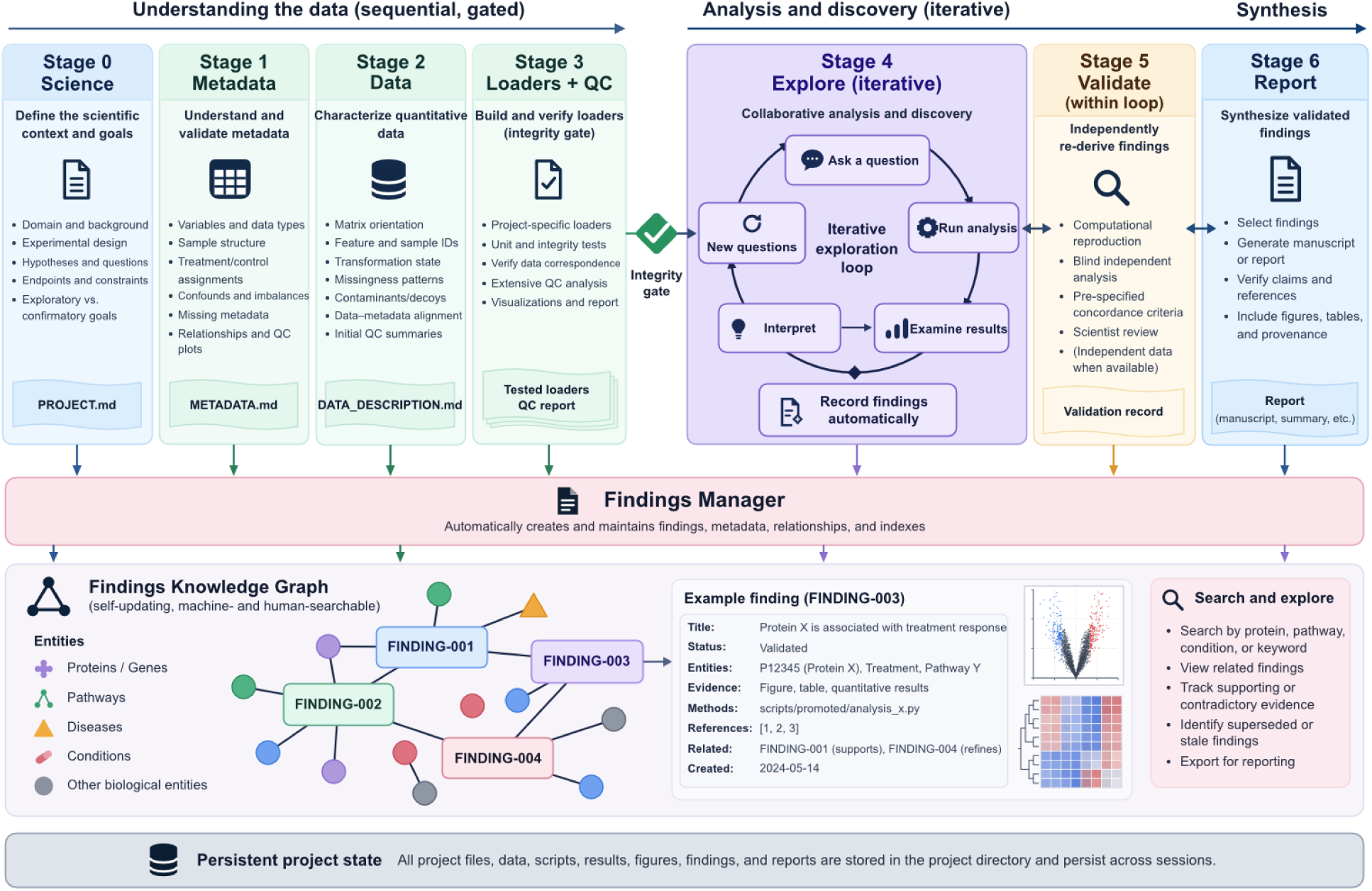
Sequential and gated analysis workflow and findings knowledgegraph. Stages 0 to 3 sequentially build on the foundations of data analysis, starting with first understanding the project and its science, then understanding the metadata, then the data, then finally building robust, tested data loaders and performing quality control (QC). Each of these steps depends on finishing the previous step. Stage 4 supports iterative, conversational, researcher-directed analysis only after stages 0 to 3 are complete, ensuring analysis only takes place in a context where the underlying data are understood and trusted.

Once the researcher is satisfied, they may validate their findings in stage 5 and, once validated, move to incorporating those findings in a comprehensive report (stage 6). All the while, substantive discoveries and observations are automatically saved by the findings-manager agent in the findings knowledge graph. Users may query the knowledge graph at any time by asking Claude and view findings files directly with any viewer capable of displaying Markdown.

#### Project Instructions and Persistent Workflow State

The Findings Workflow includes a special project `CLAUDE.md` file that is copied into a project’s directory when the plugin’s `init` command is run. A `CLAUDE.md` file is a special file that is automatically read by a Claude Code session in a given directory, in this case, the project’s directory. It supplies persistent instructions to the main Claude Code session whenever the project is opened. For the Findings Workflow, rules such as workflow ordering, use of findings, responsibilities of the main orchestrator, and human checkpoints do not depend on the researcher using the same chat session. The researcher can return days, weeks, or months later and resume the analysis using a new Claude Code session from its persisted project state without relying on the previous conversation.

#### Specialist Subagents

When the researcher launches Claude Code, by default they are interacting with the “main” agent or the “orchestrating” agent. This agent automatically reads the `CLAUDE.md` file, is onboarded to the project state, and is the agent through which the researcher interfaces with the project. This main agent is primarily responsible for orchestration: it determines what needs to be done, in what order, and dispatches the work to specialist subagents, responsible for carrying out specific tasks or managing specific kinds of data in the project. (Figure 1B) The Findings Workflow ships with 13 subagents, assigned to seven subsystems. (Table 1). These subagents run in isolated contexts. Research activity, code generation, statistical analysis, and figure construction do not continuously accumulate in the main agent’s context and create useful separations between tasks. Importantly, all the subsystems emphasize verification and review, where all claims are verified and all output is reviewed by independent subagents.

#### Reusable Skills

The Findings Workflow ships with seven reusable skills. Skills are distinct from subagents, skills define a reusable procedure for performing a particular type of work, and may be used by any agent. The seven skills included with the plugin are: (1) `verification-task-builder`, (2) `statistical-analysis`, (3) `figure-generation`, (4) `research-publications`, (5) `research-protein`, (6) `research-source-code`, and (7) `report-writing`.

#### Deterministic Enforcement Hooks

The Findings Workflow does not depend entirely on Claude following written instructions. Or put another way, not all of the workflow’s rules are prompts. The workflow includes three hooks that act as guards of data quality and integrity. These include: (1) `guard_readonly_data.py`: This prevents any modification of data in the data/directory. (2) `guard_findings.py`: This enforces strict rules about findings, figure provenance, figure and legend correspondence, linking between findings, and a structured figure hierarchy. (3) `guard_promotion.py`: This prevents analysis scripts from being promoted unless required linting and strict type checking pass.

It’s important to differentiate hooks from the concept of “gates” that exist in the Findings Workflow. A hook is a script that Claude Code runs automatically when certain events occur, such as before or after the agent uses a tool or when the user submits a prompt. Hooks are absolute and enforced programmatically, whereas gates exist as parts of prompts to the stage controller commands and may be more nuanced. For example, performing analysis of the data requires the gate in the `stage4-explore` command be passed, which requires the data to be fully understood during stage 3 first. However, knowing which user request or tool invocation actually constitutes data exploration of a type that would violate this gate is not cleanly decidable by a simple python script–it is a more complex and nuanced judgment call that is made by the main orchestrator agent.

#### Conventions and Templates for Analysis and Visualization

The Findings Workflow includes 11 conventions documents including: `findings`, `manifest`, `script-registry`, `research-corpus`, `workflow-state`, `coding`, `statistics`, `correctness`, `visualization`, `reporting`, and the `enforcement-map`. These documents define the standards against which agents and hooks evaluate work, including: statistical reporting and multiple hypothesis testing requirements, prevention of leakage during cross-validation, normalization and batch-correction practices, data-loader verification, code testing (unit tests, typing, linting, seeding, failure modes), figure formatting, color consistency, provenance enforcement, stage transitions, script promotion, and reporting rules. The enforcement-map maps individual rules to one or more concrete enforcement mechanisms (e.g., hooks, reviewer agent, orchestrator behavior, or human decision). This implements the design philosophy that a convention is only real if something actually checks it.

As of this writing, the Findings Workflow also ships with 28 registered analysis, data-processing, visualization, and support templates. These are Python implementations that have been rigorously vetted, typed, linted, tested, and designed around workflow conventions. Importantly, these templates are not meant to be used verbatim, but serve as a source of opinion downstream when researchers are using the workflow. Their orchestrator agent has access to a vetted source of opinion about data handling and preprocessing (e.g., loading data, normalization, and batch correction), statistical analysis (e.g., differential abundance using moderated linear models (limma^7^-style analysis), ordinary least squares (OLS), building classifiers, regression, enrichment analysis, and so on) and data visualization (e.g., PCA, run-to-run variance, correlation, abundance, results of feature selection, classification, regression, and enrichment). The orchestrator agent decides which template scripts are applicable to a task at hand. When used, these templates are copied into the project and adapted to the user’s specific dataset and domain. The adapted local script is recorded in the provenance of findings. And, because the script was a copy of the template, subsequent plugin updates will not alter this script or affect the provenance and reproducibility of findings. If no templates are applicable to the task at hand, the orchestrator may choose to instantiate a statistician subagent to create analysis scripts from scratch, but may use patterns present in existing template examples to drive the design of those scripts.

### Automated Findings and the Findings Knowledge Graph

At the heart of the Findings Workflow is the automatic creation and maintenance of a persistent findings knowledgebase (Figure 2). As important observations emerge during analysis, they are automatically saved as separate, uniquely identified findings rather than being lost in long conversation histories. In this way, the Findings Workflow works as a self-updating scientific lab notebook: it continuously captures what was learned, how it was learned, the supporting evidence, how to reproduce it, and how it relates to other things that were learned.

Each finding is stored in a standalone Markdown document that is designed to be both machine- and human-readable. The beginning of the file contains YAML frontmatter that contains highly structured annotations for the finding. This includes the finding status (conjecture, confirmed, rejected, etc), controlled vocabulary for biological annotation, relationships to other findings, quantitative evidence, data and script provenance, figures, and references. The human-readable portion provides a descriptive writeup of the finding. This includes an abstract, description, methods, discussion, conclusion, references, figures, tables, provenance, related findings, and other supporting information. Each figure is treated as evidence and contains its own provenance identifying code and data that produced them.

Findings are connected to each other using typed relationships such as “supports”, “refines”, “contradicts”, “supersedes”, and “closes”. They can also reference entities using controlled vocabularies, such as proteins, genes, pathways, and diseases. Together, these relationships form an always-evolving findings graph and knowledge graph that can be searched by humans or agents. For example, the agent can retrieve all findings involving a particular protein, a particular co-variate from the metadata, or a specific treatment. Agents can find all findings that may already provide evidence for or against a hypothesis and build on those. All findings include a section describing next steps related to that finding that agents can use to suggest next steps to the researcher.

A dedicated findings-manager subagent maintains the knowledgebase. It creates and updates findings, maintains relationships and indexes, records validation outcomes, and checks for consistency or staleness when underlying data or analysis code change. Directives arrive from the main orchestrator agent about which findings should be recorded or updated, and the findings-manager carries that out. The findings-manager is purely custodial, it does not make decisions about the importance or truthfulness of a result. Those decisions are the jobs of other, dedicated subagents.

### Example Usage

The Findings Workflow was applied to data associated with Wu *et al*. (2025)^8^. In this work, Wu et al. describe Mag-Net, a magnetic-bead-based workflow for enriching extracellular vesicle (EV) associated material from plasma to improve proteomic depth and reduce interference from highly abundant plasma proteins. The authors demonstrated the effectiveness of the method by performing data independent mass spectrometry (DIA) analysis of EV-enriched plasma from 40 subjects, 10 classified as Alzheimer’s disease (ADD), 10 as Parkinson’s disease (PDD), 10 as Parkinson’s disease but cognitively normal (PDCN), and 10 healthy controls (HCN). They developed a linear support vector machine (SVM)^9^ classifier that could strongly separate ADD samples from non-ADD samples.

To demonstrate the workflow, the metadata and final matrix of normalized and batch-corrected protein abundances were downloaded from the Mag-Net manuscript GitHub repository. The Findings Workflow is capable of guiding users through both normalization and batch correction; however, in the interest of brevity these pre-processed data were used. The data were placed in the “`data`” folder of a new directory and the Findings Workflow (version 0.5.1) was run on the command line using “`claude --model claude-opus-4-8`”, which specifies the Claude Opus 4.8 model. The unfiltered chat transcript and the resulting working directory, complete with findings, figures, all analysis source code, Mag-Net protein abundances and metadata, and all other files may be found at https://github.com/uw-maccosslab/manuscript-findings-workflow. Also in the interest of brevity, the workflow was run with relatively simple goals: build a classifier to distinguish ADD from the other sample types, find proteins that are important for distinguishing these classes, and characterize those proteins. This summary of the chat session is not comprehensive, please review the GitHub link above to review all findings and figures.

#### Project Setup and Scaffolding

The Findings Workflow was installed according to the instructions at https://github.com/mriffle/findings-ai-collab-workflow, consisting of two simple commands typed into Claude Code. Afterwards, the first command given to the workflow was “`/findings-workflow:init`”, which caused Claude Code to scaffold the project. This creates directories and empty versions of files used to track project information and state. The next command was “`/findings-workflow:setup-env`”, which caused Claude Code to ensure a suitable python environment was set up. No further input was required by the user for either step.

#### Stage 0: State the Science

The next command was “`/findings-workflow:stage0-science`”. This initiates an interview of the scientist by the workflow regarding their project. Claude Code specifically responded:

“I’ll capture the scientific framing before we touch any data. This is an interview — one question at a time — and your answers shape what I ask next. Let me start broad: What is the scientific domain and system here? Tell me about the field, the organism or sample type, and the measurement technology. For instance — is this a mass-spec proteomics study, and if so, on what kind of samples (cell lines, tissue, plasma…) and from what organism? Enough context that I could correctly interpret a finding later.”

After the first response, Claude asked two more clarifying questions about the study, announced it was ready, showed a summary, wrote the project description to the “PROJECT.md” state file, and suggested moving on to the next stage.

#### Stage 1: Understand the Metadata

The next command given was “`/findings-workflow:stage1-metadata`”. This causes Claude to identify which file contains the metadata, understand its format, check its format and validity, to interrogate and understand the meaning of the columns, characterize the cohorts, look for confounders and class imbalances, and to test relationships between columns the column definitions imply should be true. In this example, Claude identified “cohort” as a possible confounder for our group of interest, the “ADD” group, as all instances of ADD subjects came from the same cohort site (Figure 3).

**Figure 3.**
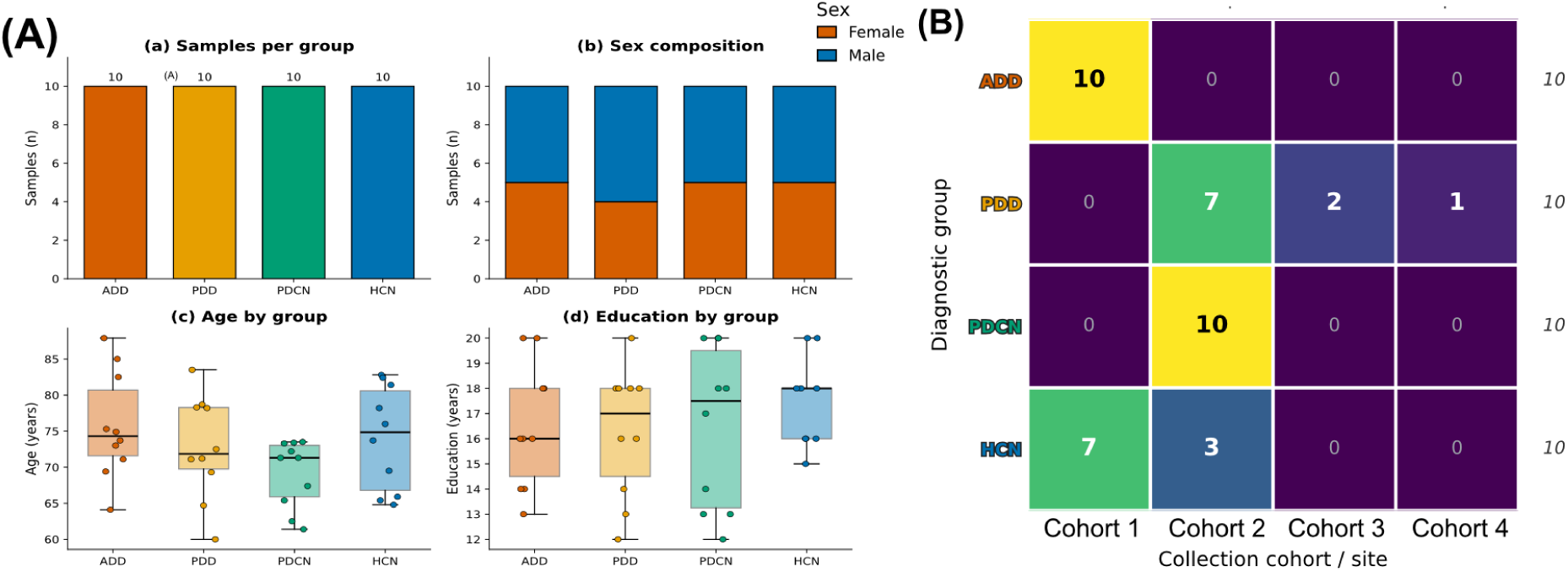
Select examples of metadata figures output by the Findings Workflow for the example analysis. (A) The cohort characterization. The top left panel shows the number of subjects for each disease state. The top right shows the distribution of sex by disease state. The bottom left shows the distribution of subject ages by disease state. The bottom right shows the distribution of education level by disease state. (B) ADD diagnosis is confounded by cohort site. The confusion matrix displays the number of subjects with each combination of disease state and cohort site.

Claude confirmed its interpretation of the metadata columns with the scientist, dispatched the findings-manager to record the confounding observation, dispatched figure generation and review subagents to create metadata figures (Figure 3), updated the “METADATA.md” state document, and suggested moving on to the next stage.

#### Stage 2: Understand the Data

The next command given was “`/findings-workflow:stage2-data`”. Similar to the metadata stage, Claude will now examine the data matrix (or data matrices) that contain the experimental results to be analyzed. For this proteomics dataset, it determined the matrix orientation, transformation (e.g., log2) state, normalization state, characterized contaminants, identified decoys, and identified duplicate data. After examining the data, Claude noted the correct orientation, that the data were log2 transformed, and already median normalized. It inquired whether batch correction should be applied, whether the data should be further normalized, that two proteins (IMPA3 and CD048) were missing values in many samples, and that yeast enolase was found in the sample and whether it should be considered for downstream analysis. Claude was told to exclude yeast enolase from analysis, to ignore the two largely missing proteins, not to re-normalize the data, and that the data were already batch corrected. Claude then performed an analysis to verify the claim the data were already batch corrected, recorded this as a finding, summarized the details about the data, recorded the final data information in the “`DATA_DESCRIPTION.md`” state file, and suggested moving to the next stage.

#### Stage 3: Build the Data Loaders and Run Quality Control Report

The next command given was “/`findings-workflow:stage3-loaders`”. This stage of the workflow carries the largest load and has two primary goals: (1) building the official data loaders that will be used for all downstream analysis and (2) performing a quality control (QC) analysis of the data, generating QC figures, and generating a QC report. Claude first worked to develop the official data loaders. It reviewed its guidance and plugin-encoded contracts then dispatched coding agents to build the loaders. This process writes the code for loading data and metadata and rigorously tests that the data are being loaded in correctly and that the mapping between the data and metadata is correct. Claude dispatched independent code review agents to review the code and correctness and appropriateness of the tests. Once all code has been reviewed and verified, Claude moves on to the QC analysis, which is built on top of the verified data loaders.

Claude loaded the data and verified aspects of the data, such as class imbalances, that were previously recorded. Once verified, it dispatched multiple figure generation subagents to generate QC figures, including PCA plots, correlation plots, abundance distribution plots, coefficients of variation (CV) plots, and dynamic range plots (Figure 4). Each of these ran in parallel, and each were subject to independent review and fix cycles until they were considered complete and correct.

**Figure 4.**
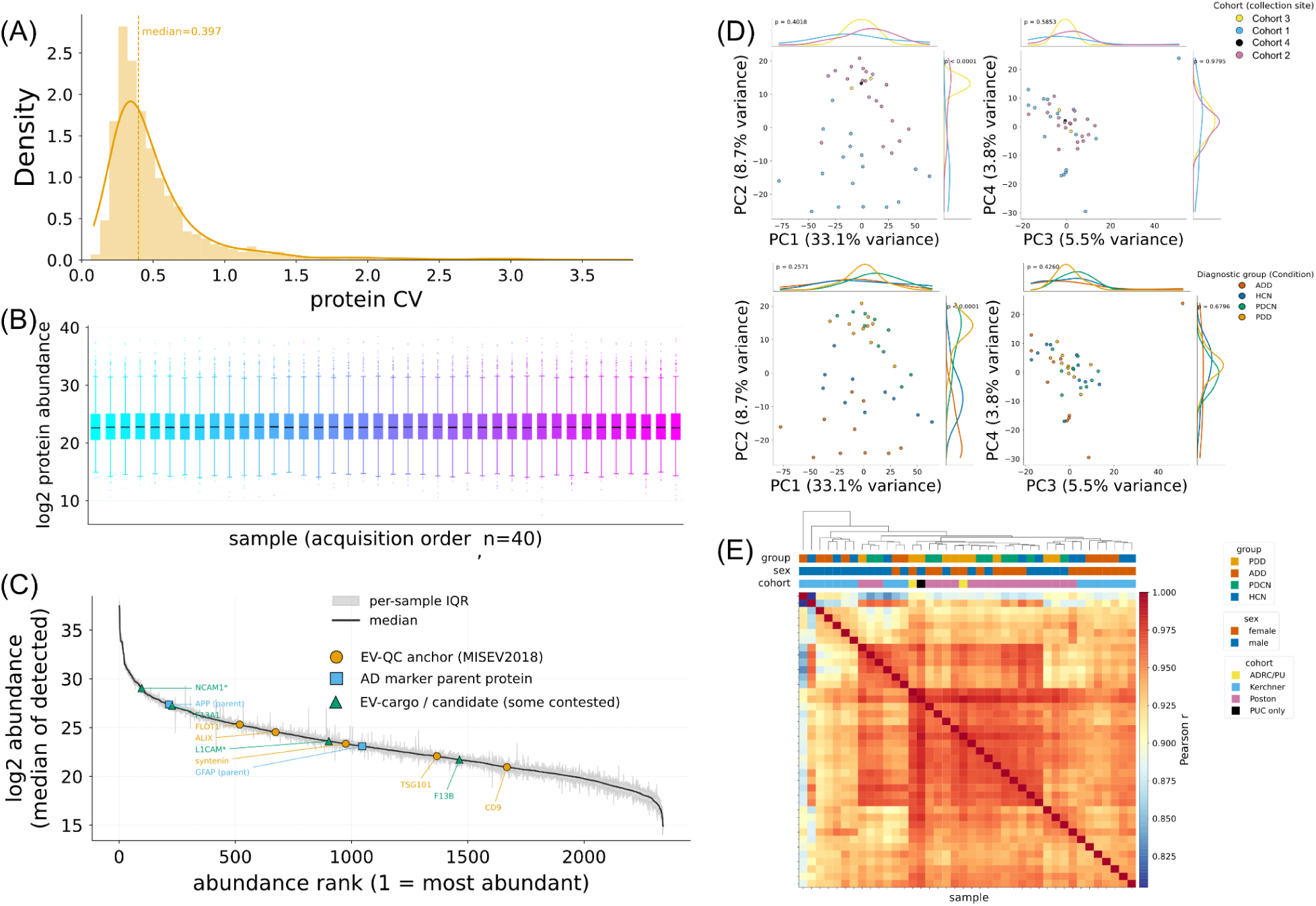
Examples of QC plots generated by the Findings Workflow applied to the Wu *et al*. dataset. (A) The distribution of the protein coefficients of variation (CV) across all samples. (B) The distribution of all measured protein abundances across all samples. (C) A dynamic range plot, where proteins were sorted by their median abundance and plotted as their abundance versus their rank. The black line notes their median abundance, and the gray bars note the interquartile range for each protein’s abundance across all samples. Individual proteins of interest found by the librarian subagent for EV markers of Alzheimer’s disease are noted, with shapes and color indicating classifications of those proteins determined by the librarian agent. (D) Principal Component Analysis (PCA) plots, colored by cohort (top) and diagnostic group (bottom). The marginal plots indicate the density of samples with a given color across each of the four principal components, and the p-value is a Benjamini-Hochberg^10^ adjusted p-value from the Kruskal–Wallis test, testing whether the observed differences among group distributions are greater than expected if the groups came from the same distribution. (E) A correlation plot examining the correlation of all samples versus all others. The samples are hierarchically clustered, with the dendrogram visible at the top. The cells are colored according to sample to sample Pearson correlation. Three colored bars at the top indicate sample labels for diagnostic group, sex, and cohort.

During the process, the scientist also asked, “Are there known plasma EV markers for Alzheimer’s we can show on the dynamic range plot?” Claude responded by dispatching the librarian agent, which in turn launched researcher subagents to query the literature and other internet sources for these markers. The findings of all the researcher subagents were verified using independent reviewer subagents, and some were sent back to the researchers for corrections, until final results were accepted. The librarian agent collated the results, wrote to its research knowledge graph, and returned that information to the main agent. The main agent then dispatched a new figure subagent to update the dynamic range plot to note the positions of the researched, known EV markers for Alzheimer’s. This figure was then reviewed, and once verified as correct, saved (Figure 4C). From this point forward, the discovered EV markers were available for downstream analysis without further research being necessary. All discovered markers are annotated with the appropriate references.

Once the QC figures were generated, reviewed, and verified, Claude dispatched subagents to generate and verify the QC report, and dispatched the findings manager subagent to record substantive findings generated during the stage 3 process. Once finished, Claude announced, “Stage 3 — Integrity gate PASSED”, summarized the results from stage 3, and suggested moving on to Stage 4 (providing the exact syntax) to begin exploring and analyzing the data.

#### Stage 4: Explore and Analyze the Data

The next command given was “`/findings-workflow:stage4-explore`”. This moves the state of the project to the data analysis and exploratory stage, where the scientist and Claude begin collaborating on data analysis and findings generation. First, Claude verified that all of stages 0 through 3 were complete. It then stated that since distinguishing the ADD group from all others was our primary aim, it dispatched a statistician subagent to begin performing differential abundance analysis to yield interpretable per-protein hits that will inform related steps, downstream. After the statistician agent finished and verified, Claude reported a summary of the results. It found 363 proteins statistically significantly differentially expressed with a Benjamini-Hochberg (BH) q-value less than 0.05 (133 up and 230 down in ADD). It warned that the shape of the distribution of the p-values was suspicious and that the count of significant proteins may be inflated. And again reminded us of a possible cohort confound discovered earlier and advised caution in interpreting the results. It also noted that none of the known plasma EV ADD markers discovered by the librarian previously were found to be significant. Claude then dispatched parallel figure generation subagents to produce and verify a Volcano plot (Figure 5A) and p-value histogram figure.

**Figure 5.**
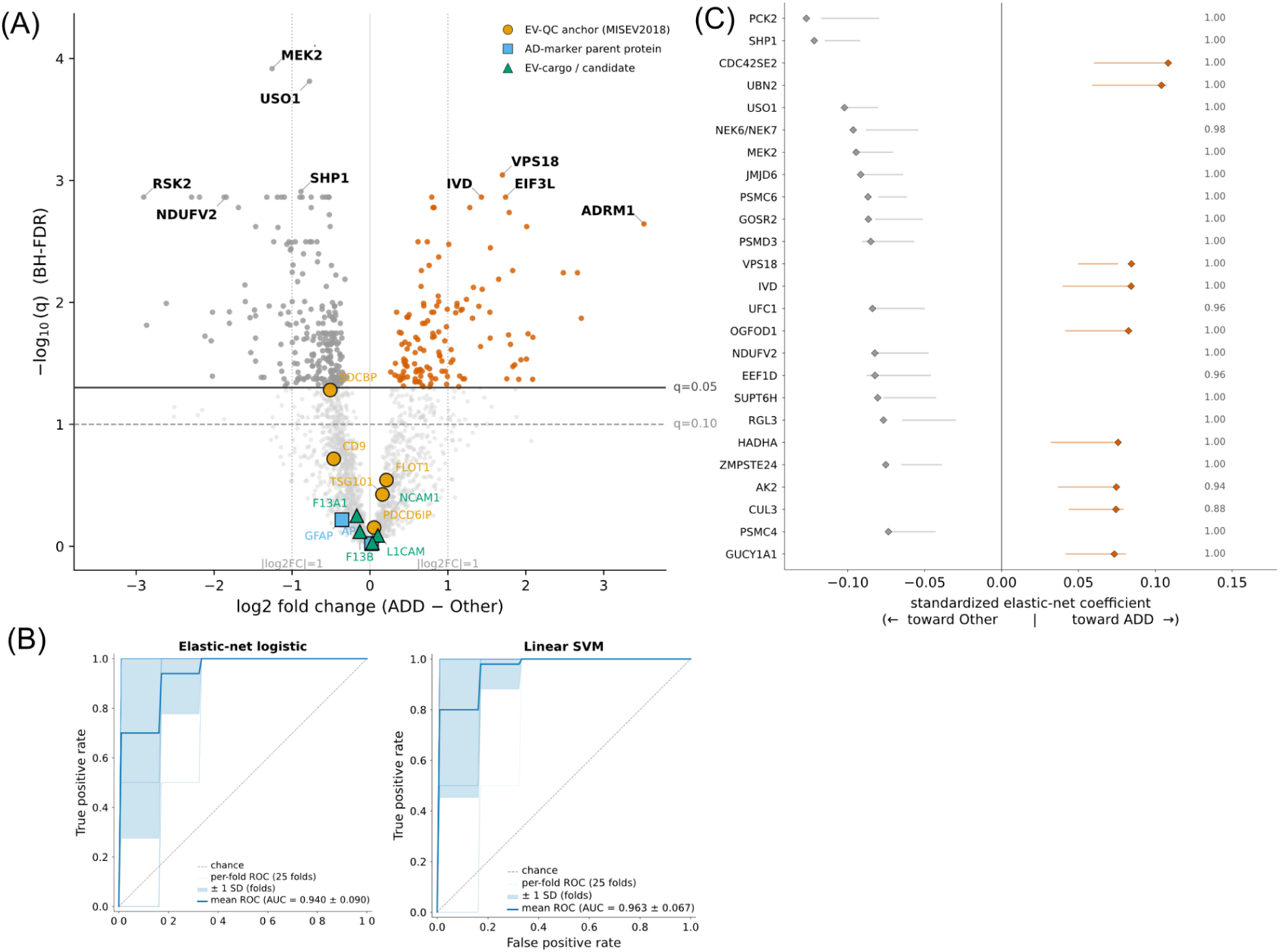
Select figures generated by the Findings Workflow for differential abundance analysis and classifier modelling in the Wu *et al*. dataset. (A) Volcano plot of differential protein abundance between ADD and the pooled PDD, PDCN and HCN groups, estimated with a moderated (empirical Bayes^11^) linear model. Each point is one protein. The horizontal axis is the log2 fold change (ADD minus pooled groups) and the vertical axis is the −log10 Benjamini–Hochberg adjusted q-value. The solid horizontal line marks q = 0.05 and the dashed line q = 0.10. Dotted vertical lines mark a two-fold change. Proteins with q < 0.05 are colored by direction (orange, higher in ADD; grey, lower in ADD), and proteins with q ≥ 0.05 are shown in light grey. Previously reported EV and AD reference proteins are overlaid as larger, black-outlined symbols and labeled with their gene names; symbol shape and color indicate the class. (B) Receiver operating characteristic (ROC) curves for classifying ADD versus the pooled PDD, PDCN and HCN groups using all measured proteins, with an elastic-net logistic regression^12^ (left) and a linear SVM (right). Performance was estimated by nested cross-validation (stratified 5-fold, repeated 5 times; hyperparameters tuned within the inner loop), and each ROC curve is computed only from samples held out from training. Thin translucent lines are the 25 individual outer-fold ROC curves. The bold line is the mean ROC curve across folds and the shaded band is ±1 standard deviation. The diagonal line marks chance performance. (C) Elastic-net logistic regression coefficients for classifying ADD versus the pooled PDD, PDCN and HCN groups, showing the 25 proteins with the largest absolute coefficients. Diamonds mark each protein’s coefficient in the final model fit to all samples; positive coefficients (orange) push predictions toward ADD and negative coefficients (grey) toward the other groups, with the vertical line at zero. Horizontal bars show the interquartile range of the coefficient across repeated refits on cross-validation training subsets at the same hyperparameters, computed over the refits in which the protein was selected. Numbers at the right give each protein’s selection frequency, the fraction of refits in which its coefficient was non-zero.

Based on the scientist’s stated goals in Stage 0, Claude suggested building a logistic regression classifier with elastic net^12^ regularization. The scientist asked why this particular classifier was chosen, and Claude responded with the reasoning behind the choice and suggested alternatives such as a linear support vector machine (SVM), along with the reasoning for those suggestions. The user responded that they’d also like to explore a linear SVM. Claude then dispatched statistician subagents to build, run, and verify the classification analyses. The classifier code was based on template classifier code included with the plugin, but tailored for this analysis as Claude judged appropriate. Evaluation of the classifiers was done using nested 5-fold cross-validation repeated 5 times, to ensure no data leakage occurred, even when choosing hyperparameters. After the classifiers ran, Claude reported back with initial results. The logistic regression classifier distinguishing ADD samples from all others reported a ROC-AUC of 0.94 and balanced accuracy of 0.83. The linear SVM classifier reported a ROC-AUC of 0.96 and a balanced accuracy of 0.74. The SVM classifier ROC-AUC was nearly identical to that reported in Wu *et al.*, which built the same classifier using a linear SVM. It stated that both methods essentially tied and there was no clear winner. It reported that the top features used by the classifiers were stable and coherent. The top 25 logistic regression features were selected in 100% of the cross-validation folds and 23/25 were significant differential abundance features. It then dispatched subagents for the findings manager to record the finding and figure generation agents to build and verify ROC figures (Figure 5B) and logistic regression feature stability figures (Figure 5D).

Claude then initiated a conversation about where to take the analysis next. Claude suggested running a label-shuffle null for the classifiers to establish the statistical significance of the observed ROC AUCs, but noted this still would not address the potential issue of the cohort confound. So it also suggested building a classifier to predict cohort 1 vs cohort 2 among the 30 non-ADD samples (HCN and PDD span both sites). If the proteome predicts the cohort at a similar AUC as the ADD classifiers, that is evidence that the ADD signal is substantially just predicting the site. The scientist elected to perform both analyses. Claude dispatched statistician subagents to build, perform, and verify the analyses. This reported back a null mean ROC AUC of 0.5 for both the logistic regression and linear SVM classifier, with 95% confidence intervals of 0.24 to 0.78 and 0.23 to 0.76, respectively. The reported p-value for the observed ROC AUC on unshuffled data was reported to be less than or equal to 0.001. The site probe (predicting cohort 1 vs cohort 2 among the 30 non-ADD samples) reported a classifier no better than chance in predicting the site, providing evidence against the ADD classifier only predicting the site. Claude reported the bottom line at this point as, “there’s a genuine, reproducible, null-beating signal that distinguishes ADD — stable, agreeing with the DE, sitting in intracellular proteins mostly lower in ADD,” but added that the ADD samples were still fundamentally confounded with site. Claude then dispatched the findings manager to record the findings and figure generation agents to generate the figures.

The scientist was then interested in performing an enrichment analysis of significant features. Claude asked the user which set of features to use for the analysis, and suggested running a Boruta^13^ analysis to find features. Boruta is an all-relevant feature-selection method built around Random Forests^14^ that compares the importance of each observed feature against randomized “shadow” versions of the features, retaining those that consistently carry more predictive information than expected by chance. This approach is well suited to high-dimensional proteomics data because it can identify multiple potentially informative proteins without requiring linear relationships or selecting only a minimal predictive subset. The scientist noted that Boruta wouldn’t indicate which features are enriched or depleted, and Claude suggested running Boruta and labelling the features using their sign from the previous differential abundance analysis. Then, enrichment analysis would be performed separately on the Boruta “up” features and “down” features. The user agreed, and Claude dispatched a statistician subagent to run and verify the Boruta analysis. Boruta found 40 confirmed features, 18 up and 22 down, in ADD. 33/40 were in the logistic regression feature set. 7 were labeled as “redundant-but-real”, where they were confirmed as real by Boruta but were dropped by regularization during logistic regression, which is exactly what Boruta is meant to correct for. 38/40 were also differential abundance hits. Claude dispatched the findings manager and figure subagents to build related findings and figures (Figure 6A). Claude then dispatched a statistician subagent to perform an enrichment analysis using g:Profiler^15^, which uses a cumulative hypergeometric test to determine whether proteins annotated with a given Gene Ontology (GO)^16^ or KEGG^17^ term are represented in the observed protein list more frequently than expected by chance relative to the background set, with multiple-testing correction performed using g:Profilers’s g:SCS method. For this analysis, the background was the set of all measured proteins. This analysis produced zero significant terms in either the up or down direction. Claude dispatched findings and figure agents to note the finding.

**Figure 6.**
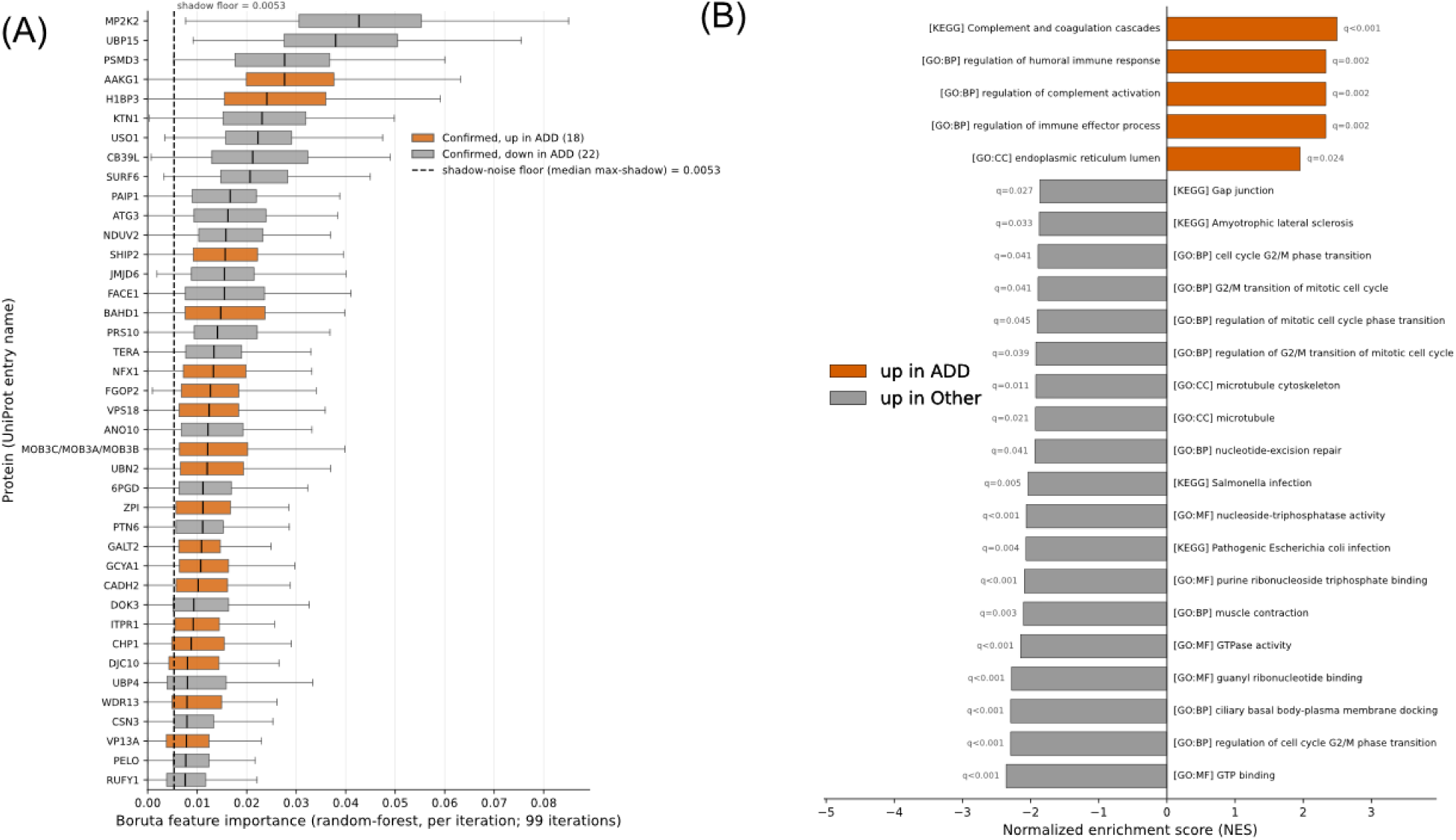
Figures generated by the Findings Workflow for Boruta and GSEA^18^ enrichment analysis from the Wu *et al.* dataset. (A) The 40 proteins confirmed by Boruta all-relevant feature selection as distinguishing ADD from pooled PDD, PDCN and HCN. The horizontal axis is Boruta feature importance: the random-forest importance of each protein, recorded at each of 99 Boruta iterations. Proteins are ordered on the vertical axis by median importance. Each box summarizes one protein’s importance across iterations (box, interquartile range; center line, median; whiskers, 1.5× IQR; outliers not shown). Orange boxes are proteins with higher abundance in ADD, and grey boxes lower abundance, based on the direction of change in the differential abundance analysis. The dashed vertical line is the shadow-noise floor: the median, across iterations, of the highest importance reached by any shadow (randomly shuffled) feature. (B) Gene set enrichment analysis (GSEA) of ADD vs. pooled PDD, PDCN and HCN (n = 10 vs. 30). All measured proteins (n = 2,324) were ranked by moderated t-statistic, and GO and KEGG gene sets were tested for clustering at either end of the ranking. Shown are the 24 gene sets with the largest absolute normalized enrichment score (NES) among the 39 significant at FDR q < 0.05 (q-values at bar ends). Bar length is the NES, which measures how strongly a set’s members cluster at one end of the ranked list. Orange sets (positive NES) are increased in ADD; grey sets (negative NES) are decreased in ADD.

Here the user pointed out that the enrichment analysis had only tested the number of distinct proteins annotated to each GO or KEGG term, not their abundance, so a pathway could still be over-represented when abundance was taken into account. Claude agreed and identified a rank-based gene set enrichment analysis (GSEA)^18^ as the appropriate test. Claude asked which statistic to rank proteins by, recommending the moderated t-statistic^11^, which the user accepted. Claude spawned a statistician subagent to rank all 2,340 proteins by moderated t, map UniProt^19^ accessions to HGNC^20^ symbols using MyGene.info^21^ (2,324 mapped), and run gseapy^22^ prerank against the Enrichr^23^ 2021 Gene Ontology biological process, molecular function and cellular component libraries and KEGG, with 1,000 permutations. The analysis produced substantive results (39 gene sets at FDR < 0.05), including a complement/coagulation and humoral-immune secretory program increased in ADD and GTPase-related sets increased in the pooled non-ADD groups. Claude dispatched figure-generator agents to produce the figures, each checked by an independent figure-reviewer agent, and then the findings-manager agent to record the result as a finding (Figure 6B).

The 39 significant gene sets identified by GSEA collapse to about six themes. Proteins lower in ADD were dominated by small GTPases and microtubule-based transport proteins, in agreement with Wu et al., who noted that proteins with GTPase activity and microtubule-transport proteins, including kinesins and dynactin complex members, were decreased in ADD relative to the other three groups; our differential abundance analysis also reproduced the direction of all ten proteins that study highlighted as increased or decreased in ADD, each at q < 0.05. Proteins higher in ADD were dominated by complement/coagulation proteins and serine-protease inhibitors, a pattern not described in the original report. It is consistent with higher abundance of several complement- and coagulation-related proteins, including serine-protease inhibitors, in plasma-derived extracellular vesicles from patients with AD^24^, altered plasma complement protein levels^25^, elevated CSF clusterin and C3 in AD^26^, complement activation products in AD brain^27^, and contact-system activation in AD plasma^28^. However, it’s important to note that since ADD diagnosis was confounded with cohort site in these data, we cannot exclude the possibility that the observed enrichment reflects site-associated effects rather than disease biology.

#### Stages 5 and 6: Findings Validation and Report Generation

Although not described here in the interest of brevity, the user could have triggered an independent validation of any finding by running a command such as “`/findings-workflow:stage5-validate 7`” or conversationally asking to validate the specific finding. This takes a “candidate” finding recorded during a previous stage and attempts to move it towards “validated” by having a subagent attempt to independently derive the result without seeing the answer. For example, the main orchestrator may ask the subagent, “What proteins are enriched in the ADD samples?”. It would never ask the subagent to confirm that a specific list of proteins were enriched in the ADD samples. Additionally, the user could have triggered the generation of a comprehensive report by running “`/findings-workflow:stage6-report`”. This builds a report from the evidence, figures, methods, caveats, references, and status of each finding. The Findings Workflow will ask the user which mode to run in: the “qc” option builds a data quality report and the “research” option (the main deliverable) selects validated findings to build a story to aid in manuscript preparation. The final report includes a title, abstract, methods (derived from findings), results (including figures and tables), discussion, and references. Each section is built by a writer and report-reviewer subagent pair, with iterations on writing and reviewing until each section passes review. Reviewers work to ensure each claim has a source and that each sentence in results and discussion must trace to a finding or vetted research finding. A final reviewer stitches the sections together, removes repetition, fixes inconsistent terms, and checks cross-references. The report is generated as Markdown and PDF documents suitable for sharing. This is not intended to generate manuscripts for publication, only to collate findings and aid in downstream interpretation and sharing of complex results.

## Discussion

### Evaluation

A LLM-backed workflow like the Findings Workflow is difficult to objectively evaluate. Questions regarding whether or not using the workflow leads to better results are certainly legitimate, though what constitutes a “better result” is open for interpretation. The authors have received very positive and enthusiastic informal feedback from test users, but fully vetting the impact of the workflow on research in a statistically meaningful way is beyond the scope of this manuscript. Additionally, it is also likely that two different users using the same workflow to analyze the same data with the same interests could conceivably end up with slightly different analyses, but likely the same outcomes. These differences would be driven by the exact syntax researchers used to state their project goals, metadata descriptions, or prompts given during data analysis. The authors believe this to be not dissimilar from two researchers analyzing data, themselves.

### Security and Confidentiality

As currently designed, the Findings Workflow runs within Claude Code, which sends user prompts and model outputs, including any files or data Claude reads, to the model provider over an encrypted connection. How that data is handled depends on the user’s account rather than on the workflow. On consumer plans (Free, Pro, Max), Anthropic trains on user data if the model-improvement setting is on, retaining it for five years, or 30 days if the setting is off. Under commercial terms (Team, Enterprise, API, and third-party cloud platforms), Anthropic does not train on this data by default and retains it for 30 days. Organizations may have stronger arrangements, such as zero data retention, a Business Associate Agreement for health data (which covers Claude Code only with zero data retention enabled), or access through Amazon Bedrock or Google Cloud under that provider’s terms. Users handling confidential data should confirm which terms apply to their account, whether their institution holds such agreements, and whether requirements such as HIPAA, FERPA, or IRB protocols impose additional conditions.

### Practical Considerations and Token Cost

A token is a small chunk of text that an LLM model reads or writes. Subscriptions to LLM services, such as Claude, come with a quota of tokens a user may consume during a period of time, such as within a 5-hour or 7-day window. Using the Findings Workflow can consume a large amount of available tokens, particularly when performing a large amount of research. As of this writing, the authors recommend using Opus 5.5 for the orchestrating agent (main interface). If on the $20/month tier, the authors recommend using Sonnet 5.5 for your orchestrating agent. This will lead to substantially lower token usage, but may lead to less optimal output. Note that as of this writing, the research, coding, and figure generation subagents are configured to use the Sonnet model to help reduce cost, regardless of whether or not your orchestrating agent is set to Opus.

### Workflow Agnostic

Because the Findings Workflow operates on the end results of quantitative analysis pipelines, it is workflow agnostic and not tied to any specific quantitative analysis software or -omics workflow. Beyond DIA proteomics, the workflow has been applied successfully to targeted proteomics methods such as parallel reaction monitoring (PRM), data-dependent acquisition (DDA) MS1-based label free quantification (LFQ), and non-proteomics methods such as transcriptomics data. Although the plugin ships with quality control and analysis templates designed in the context of quantitative proteomics, the staged, gated design and templates are applicable to a very wide range of quantitative -omics data beyond proteomics. The templates are vetted implementations of quality controls, analysis, and visualization techniques that are generally applicable to quantitative data. They may be used directly in other domains, or (since the workflow is backed by an LLM) they may be customized or used as a source of truth for the design of new scripts specific to a new domain.

### The Value of Saved State

The Findings Workflow very strictly enforces the saving of state as work is performed. It saves which stage the analysis is in, which analyses have been performed, and (via the findings manager) has a record of all that has been discovered so far and how those results were derived. This strict recording of state ensures that any new Claude Code session started in the project directory is comprehensively onboarded, it knows precisely what has been done and knows what should be done next. This allows users to confidently and routinely restart Claude Code when their main orchestrator agent’s context is getting too full. It is well known that the performance of LLM agents will begin to suffer when their context starts to fill up, a phenomenon colloquially referred to as “context rot”. If the user notices this occurring, there is very little cost in simply clearing the context and starting over with a fresh agent that will perform better. Additionally, this allows users to return to the project after enough time has passed that they may have forgotten that state of the project or details about what has been done. A new agent onboarded months after the last work was performed will be immediately up to date and able to continue the analysis as if no time has passed.

### Future Directions

Work is currently underway to expand the findings workflows to other agentic harnesses. Examples include the Codex^29^ harness from OpenAI. Additionally, we have planned expansion to so-called “open” harnesses, such as OpenCode^30^ and Deepseek Harness^31^. These open harnesses could simplify the process of using open weight models as the LLM that drives the Findings Workflow. These open weight models typically cost much less than frontier models from companies such as Anthropic and OpenAI, but may not perform at the same level. However, they may perform at a sufficient level, supporting data analysis at a greatly reduced cost. This should also simplify running the LLM using locally-hosted hardware, which should address issues of privacy and on-going cost (after purchase of the initial hardware), as open weight models may be hosted locally for free. Work is also under way to expand the example templates included with the plugin to include other types of quantitative proteomics experiments (such as parallel reaction monitoring (PRM), Tandem Mass Tags (TMT), and DDA label-free quantification (LFQ). Additionally the authors are working to expand example templates beyond proteomics altogether, by including examples for transcriptomics, metabolomics, and other omics. The authors are also working to expand the templates to include other types of downstream biological interpretation scripts, especially for taxonomic and functional characterization and analysis in the context of metaproteomics and microbiome analysis.

## Conclusions

LLMs make sophisticated statistical and computational analyses more accessible to scientists who are not experts in statistics or programming. However, analysis conducted through a conversation does not ensure the underlying data are understood, that the analysis was appropriate, or that the resulting figures and conclusions can be reproduced. Because LLMs sound reasonable and confident, mistakes they make may be particularly hard for non-experts to spot. The Findings Workflow was developed to place structure and rigor around this process. It employs a staged and gated analysis workflow that ensures data are understood and loaded correctly before they are analyzed. It employs specialist subagents, independent review and validation, persistent project state, vetted analysis templates, and (critically) the automated recording of findings and their provenance. The result is an analysis environment in which the conversational flexibility is retained while the data, code, methods, figures, and conclusions produced remain accessible and reproducible.

As a simple demonstration, the Findings Workflow was applied to a published plasma EV proteomics dataset to develop a classifier, identify a set of biomarkers for Alzheimer’s disease, and characterize that protein set. The workflow identified a strong cohort-site confound before analysis began, reproduced the strong classifier performance, reproduced the reported directions of change for highlighted proteins, and independently recovered related functional signals. It also identified possible enrichment of complement, coagulation, and humoral immune processes among proteins increased in Alzheimer’s disease.

The Findings Workflow is free and open source and is available at https://github.com/mriffle/findings-ai-collab-workflow. The complete example analysis described here, including the unfiltered chat transcript, findings, figures, source code, and analysis artifacts, is available at https://github.com/uw-maccosslab/manuscript-findings-workflow.

## Acknowledgements

This work is supported by the University of Washington’s Proteomics Resource (UWPR95794). The authors would like to thank members of the MacCoss and Isoherranen labs at the University of Washington for their assistance testing and evaluating the Findings Workflow.

## Competing Interest Statement

The MacCoss Lab at the University of Washington has a sponsored research agreement with Thermo Fisher Scientific, a manufacturer of mass spectrometry instrumentation. M.J.M. is a paid consultant for Thermo Fisher Scientific. The MacCoss lab receives funding from Agilent, Bruker, Sciex, Shimadzu, Thermo Fisher, and Waters for the maintenance and support of the Proteowizard and Skyline open source software projects. The remaining authors declare no competing interests.

## Notes

https://github.com/uw-maccosslab/manuscript-findings-workflow

